# Captive rearing effects on the methylome of Atlantic salmon after oceanic migration: sex-specificity and intergenerational stability

**DOI:** 10.1101/2022.10.03.510655

**Authors:** Clare J. Venney, Raphaël Bouchard, Julien April, Eric Normandeau, Laurie Lecomte, Guillaume Côté, Louis Bernatchez

## Abstract

Captive rearing in salmon hatcheries can have considerable impacts on both fish phenotype and fitness within a single generation, even in the absence of genetic change. Evidence for hatchery-induced changes in DNA methylation is becoming abundant, though questions remain on the sex-specificity of these effects, their persistence until spawning, and potential for transmission to future generations. Here we performed whole genome methylation sequencing of fin tissue for 16 hatchery and 16 wild Atlantic salmon (*Salmo salar*) returning to spawn in the Rimouski River, Québec. We identified two cohorts of hatchery-reared salmon through methylation analysis, one of which was epigenetically similar to wild fish, suggesting that supplementation efforts may be able to minimize the epigenetic effects of hatchery rearing. We found considerable sex-specific effects of hatchery rearing, with few genomic regions being affected in both males and females. We also analysed the methylome of 32 F1 offspring from four groups (pure wild, pure hatchery origin, and reciprocal hybrids). We found that few epigenetic changes due to parental hatchery rearing persisted in the F1 offspring though the patterns of inheritance appear to be complex, involving nonadditive effects. Our results suggest that the epigenetic effects of hatchery rearing can be minimal in F0. There may also be minimal epigenetic inheritance and rapid loss of epigenetic changes associated with hatchery rearing. However, due to sex-specificity and nonadditive patterns of inheritance, methylation changes due to captive rearing are rather complex and the field would benefit from further research on minimizing the epigenetic effects of captive rearing in conservation efforts.

## Introduction

Many natural populations are in decline due to human activities (Jenkins, 2003), leading to conservation efforts to bolster populations. Certain economically or culturally important fish species are particularly threatened by human-induced changes such as habitat loss, overexploitation, and climate change (Arthington, Dulvy, Gladstone, & Winfield, 2016). Captive breeding is often used both to replenish populations in decline and to supplement populations for recreational fishing purposes (Fraser, 2008), with over 300 species of fish reared in hatcheries for supplementation efforts worldwide (Brown & Day, 2002). This often involves the capture of wild organisms, controlled breeding in hatcheries, and release of offspring into natural systems at various life stages. Salmonids have been a strong focus of supplementation efforts due to their economic and recreational value (Fraser, 2008).

Despite the high costs and ecological importance of salmonid supplementation, hatchery rearing has resulted in unintended phenotypic and fitness consequences. Hatchery-reared salmon often have altered phenotypes and reduced fitness relative to wild fish. A multispecies meta-analysis showed that hatchery fish had 50% lower reproductive success than wild fish (Christie, Ford, & Blouin, 2014) while a multidecade monitoring study in Ireland showed that hatchery-reared Atlantic salmon (*Salmo salar*) had 36% the lifetime reproductive success of wild conspecifics (O’Sullivan et al., 2020). Decreases in reproductive success due to hatchery rearing depend on life stage at release (Bouchard, Wellband, Lecomte, Bernatchez, & April, 2022; Milot, Perrier, Papillon, Dodson, & Bernatchez, 2013) and are generally more pronounced in male salmon, potentially due to reduced sexual selection and higher rates of precocious maturity in hatchery environments (Christie et al., 2014). Hatchery rearing is also associated with phenotypic changes in salmon, including reduced swimming capacity (Pedersen, Koed, & Malte, 2008), differences in heart morphology and function (Frisk et al., 2020; Leonard & McCormick, 2001), and altered gut microbiota (Lavoie, Courcelle, Redivo, & Derome, 2018). These phenotypic changes often arise after a single generation of captive rearing, prompting interest in the molecular causes of phenotypic and fitness differences.

Initially, hatchery rearing was thought to induce genetic changes, though genetic changes are often minimal and transcriptional changes are more pronounced. A single generation of hatchery rearing is generally insufficient to cause genetic differentiation between captive and wild salmon (Gavery, Nichols, Goetz, Middleton, & Swanson, 2018; Le Luyer et al., 2017), though SNPs associated with domestication have been detected after several generations of hatchery rearing in Atlantic salmon (Harder & Christie, 2022). However, the hatchery environment has been shown to induce transcriptional changes in steelhead (*Oncorhynchus mykiss*; Christie, Marine, Fox, French, & Blouin, 2016), Atlantic salmon (Frisk et al., 2020), and Coho salmon (*O. kisutch*; Leitwein et al., 2022). This led to the idea that epigenetic mechanisms controlling gene expression, such as DNA methylation, may serve as the molecular mechanisms underlying rapid phenotypic shifts and fitness declines in hatchery environments.

DNA methylation refers to the addition of methyl groups to the DNA, generally at CpG cytosines in vertebrates (Anastasiadi, Venney, Bernatchez, & Wellenreuther, 2021). Unlike genetic variation, DNA methylation can rapidly change in response to environmental cues and can influence gene expression and phenotype (reviewed in Anastasiadi et al., 2021). Methylation differences have been reported in captive fish strains reared in typical and enriched environments (Berbel-Filho et al., 2020; Venney, Wellband, & Heath, 2021) and in European sea bass (*Dicentrarchus labrax*) undergoing domestication (Anastasiadi, Piferrer, & Wittkopp, 2019). There is also a growing body of literature supporting the effects of hatchery rearing on the methylome (Gavery et al., 2019, 2018; Koch, Nuetzel, & Narum, 2022; Le Luyer et al., 2017; Leitwein et al., 2021; Rodriguez Barreto et al., 2019; Wellband, Roth, Linnansaari, Curry, & Bernatchez, 2021). A recent review on the epigenetic effects of hatchery rearing found that these methylation differences were often targeted to genes involved in stress and immune response, growth, and neural development, consistent with the added stress and environmental simplicity of hatchery environments (Koch et al., 2022). It is also possible that methylation changes associated with captive rearing could affect multiple generations as DNA methylation patterns are often heritable (Anastasiadi et al., 2021). Methylation changes in the sperm of captive-reared fish have been reported (Gavery et al., 2018; Rodriguez Barreto et al., 2019; Wellband et al., 2021) and can persist through oceanic migrations until salmon return to spawn (Leitwein et al., 2021). One study showed epigenetic inheritance due to parental captive rearing in Atlantic salmon, though this study used smolt-to-adult supplementation, an alternative rearing technique where salmon are captured as smolts and held until adulthood (Wellband et al., 2021). In this study, many somatic methylation changes present in F0 were not passed to F1, though novel methylation changes were present in F1 and were associated with phenotypic changes (Wellband et al., 2021). The evidence for epigenetic effects of hatchery rearing is growing and has potential long-term, multigenerational implications for supplemented fish stocks.

However, some knowledge gaps on the effects of captive rearing on the methylome remain. Despite clear sex-specific effects of hatchery rearing on fitness (Christie et al., 2014), it remains unclear whether hatchery-induced methylation changes also differ between the sexes. If sex-specific methylation changes occur and persist until spawning, they could be passed on to offspring. This could lead to complex patterns of inheritance in F1, particularly when reciprocal hybrids (hatchery mother x wild father or wild mother x hatchery father) occur in supplemented populations. Thus, there is a need to assess the sex-specificity of hatchery-induced methylation changes and the potential for transmission to offspring to understand the stability and complexity of the effects of captive rearing on natural populations. In this study, we assess (i) whether the epigenetic effects of early-life hatchery rearing persist until salmon return to freshwater to spawn, (ii) whether there were sex-specific effects of hatchery rearing on DNA methylation, (iii) to what extent these methylation changes were passed on to offspring, and (iv) if there were biases in the inheritance of these marks due to maternal, paternal or nonadditive effects. This study expands our knowledge of the epigenetic effects of captive rearing that may lead to fitness declines through supplementation efforts. It also underlines the importance of considering epigenetic modifications, including their stability and heritability, in conservation management decisions when assessing the impacts of supplementation efforts.

## Materials and Methods

### Parent and offspring sampling and DNA extraction

We studied the Rimouski River population of Atlantic salmon (*Salmo salar*), located on the south shore of the St Lawrence River in Québec, Canada. Stocking has been ongoing in this river from 1992 to 2021. In this system, wild-caught fish are trapped at an impassable dam each year and transported past the dam to a 21 km length of upstream spawning grounds. Some of the captured fish are used as broodstock for the supplementation effort each year and transported to the Québec Government hatchery in Tadoussac. From 2011 to 2016, the offspring of these crosses were stocked in the river as young of the year parr with their adipose fins removed for future identification.

The samples used here were obtained from a previous study on the reproductive success of wild and hatchery-reared fish, with detailed sampling information available in Bouchard et al. (2022). In 2018, a fin clip was taken from all returning adults captured and transported up the dam to their spawning grounds. Hatchery-reared adults were identified by the absence of their adipose fin. In the following summer (July-August 2019), fry were sampled in the upstream spawning area through electrofishing and preserved in 95% ethanol. DNA was extracted from parent and offspring fin tissue using a salt-based DNA extraction method (Aljanabi & Martinez, 1997). While fin is less biologically active than other tissues, its use allowed for non-lethal and minimally invasive sampling of adults returning to spawn. Due to the tissue specificity of DNA methylation (Gavery et al., 2018; Venney, Johansson, & Heath, 2016), we used the same tissue for the offspring to allow comparisons of DNA methylation between generations.

### Parentage analysis

Parentage analysis was performed in Bouchard et al. (2022). Briefly, parental sex was verified using a PCR-based genetic sex marker for Atlantic salmon (King & Stevens, 2020) to inform parentage possibilities. Parent and offspring samples were amplified at 52 microsatellite loci using multiplex PCR panels 1a and 1b from Bradbury et al. (2018) in a two-step PCR protocol (Zhan et al., 2017). The first step used a Qiagen Multiplex Master Mix to amplify each panel independently using adaptor-tagged primers (for specific PCR conditions and reagent concentrations, see Bouchard et al. (2022)). The panels were then pooled for each sample and cleaned using Quanta Bio SparQ PureMag beads. The cleaned products were used in the second PCR step which ligated barcodes to the primer adaptors to allow multiplexing of samples during sequencing. Sequencing was performed at the Institut de Biologie Intégrative et des Systèmes (IBIS) at Université Laval, Québec. Libraries were diluted to 10-12 pM and single-end sequencing was performed using an Illumina MiSeq with the MiSeq Reagent Kit V3 with 150 cycles and dual indexing to increase multiplexing capacity.

Demultiplexing of sequence data was performed with the MiSeq Sequence Analysis software. MEGASAT was used to determine allele lengths (Zhan et al., 2017) with a minimum sequence depth of 20 reads per locus per sample required to call allele sizes. Lengths were confirmed or modified based on histogram outputs. After all processing, loci were excluded if they had more than 10% missing data. Parentage analysis was performed in Cervus v3.0 (Kalinowski, Taper, & Marshall, 2007) using a 90% confidence likelihood to find the most probable mother-offspring dyad.

### Whole genome methylation sequencing

Based on parentage analysis, we selected 32 parent and 32 offspring DNA samples for whole genome bisulfite sequencing (Figure S1). For the parents, we selected 16 distinct breeding pairs that produced offspring: eight mating pairs of wild fish and eight pairs of hatchery-origin fish. We selected eight F1 offspring from four distinct groups based on parentage: (i) pure wild origin, (ii) pure hatchery origin, (iii) wild dam and hatchery-origin sire, and (iv) hatchery-origin dam and wild sire. The 16 pure strain offspring were the product of the 16 parental breeding pairs, and no offspring shared parentage. The reciprocal hybrids allowed us to determine if the effects of hatchery rearing were parent-specific; if these groups differ, they indicate that hatchery rearing affects males and females differently and these effects can be passed on to offspring. For each sample, fin DNA extractions used for microsatellite analysis were quantified, quality-checked, and sent to the Centre d’expertise et de services at Génome Québec, Montréal, Canada, where library preparation and sequencing were performed. Whole genome Methyl-Seq was performed across four sequencing lanes on an Illumina NovaSeq6000 using S4 flow cells and paired-end 150 bp chemistry with anticipated ~15X coverage for each sample.

### Methylation data processing

Methylation data processing was performed using a pipeline available at https://github.com/enormandeau/bwa-meth_pipeline. Sequence data were quality trimmed using fastp (Chen, Zhou, Chen, & Gu, 2018) to remove sequences under 100 bp and with phred scores less than 25, and to remove the first and last base of each read. Trimmed data were aligned to the North American Atlantic salmon genome (GCA_923944775.1) using bwa-meth (https://github.com/brentp/bwa-meth). Duplicates alignments were removed using Picard tools *MarkDuplicates* (https://github.com/broadinstitute/picard). MethylDackel (https://github.com/dpryan79/Methyl-Dackel) *mbias* was used to trim the beginnings and ends of reads which often result in biased methylation calls, then *extract* was used to tabulate DNA methylation for each CpG site in each individual. Paired end reads were then merged to produce bedGraph and methylKit files for further analysis. Pipeline available at https://github.com/enormandeau/bwa-meth_pipeline.

### SNP masking and coverage filtration

C/T SNPs (and A/G SNPs for the G position of CpG sites) prevent accurate methylation calling. It is not possible to determine whether a T read is due to a true unmethylated cytosine call or a thymine mutation present at that CpG site. Whole genome SNP data were obtained from an upcoming study (Lecomte et al. in prep) on two proximal populations of Atlantic salmon in the Romaine and Puyjalon Rivers, Québec, to mask C/T SNPs from our methylation data. While whole genome SNP data are not available for the Rimouski River population, these two populations are geographically and genetically close to our studied population (Figure S2) and should suffice for masking most SNPs in our dataset. Fin clip DNA was extracted for 29 Romaine River and 31 Puyjalon River Atlantic salmon. Samples were sent for whole genome sequencing at Génome Québec on an Illumina NovaSeq6000, with anticipated 16X coverage. Raw sequence data were processed using the pipeline available at https://github.com/enormandeau/wgs_sample_preparation.

Bcftools *mpileup* and *call* (Danecek et al., 2021) were used to generate genotype likelihoods from bam files with minimum mapping quality of 5. SNP calls were then filtered with bcftools *filter* and vcftools (Danecek et al., 2011) to require minimum mapping quality of 30, minimum quality scores greater than 30, minimum genotype quality over 20, minimum sequencing depth of five reads, minor allele count of two or greater, maximum two alleles, maximum 70% missing data, and minor allele frequency ≥ 0.05. SNP calling and filtering scripts are available at https://github.com/LaurieLecomte/SNP_calling_pipeline_202106. A list of C/T and A/G SNPs was exported as a bed file and bedtools (Quinlan & Hall, 2010) *intersect -v* was used to remove these SNPs from bedGraph and methylKit files.

After this, we split the methylation dataset by generation for all future filtering and analyses. BedGraph and methylKit files were quickly filtered to remove CpG sites with less than 5 or more than 100 reads, then we retained only CpG sites with sufficient coverage in at least 80% of individuals (26 of 32) for each generation. Scripts are available at https://github.com/cvenney/dss_pipeline.

### Principal component and redundancy analyses

We used principal component analysis (PCA) and redundancy analysis (RDA) for each generation to identify overall trends in whole genome methylation. MethylKit files were imported into methylKit (Akalin et al., 2012) in R (R Core Team, 2022) and united into a single data frame for each generation with only CpG sites without missing data included. PCAs were performed using the *prcomp* function to identify overall trends in the F0 dataset due to parental sex and origin (i.e., wild or hatchery), and in the F1 dataset due to maternal and paternal origin. We also performed an RDA for each dataset using vegan (Oksanen et al., 2020) to determine the significance of model terms at the whole genome level. We tested for the effects of origin, sex, and their interaction in F0 and the effect of maternal origin, paternal origin, and their interaction in F1. Model significance was tested with PERMANOVA-like tests with 1000 iterations and the significance of individual terms was also determined for each analysis.

### Coverage filtration and differential methylation analysis

DSS (Feng & Wu, 2019) was used to smooth methylation data over 500 bp regions and implement models to test for differentially methylated loci (DMLs, i.e., CpG sites) and differentially methylated regions (DMRs, identified by a high frequency of DMLs within a region). For all analyses, DMLs were considered significant if they had an FDR-adjusted p-value less than 0.05. DMRs were called based on DML p-values before FDR adjustment, as recommended by the developers of DSS, which may lead to the identification of fewer DMLs than DMRs. We further filtered the DMR results, requiring a minimum length of 100 bp, at least 10 CpGs with at least 50% of them identified as DMLs (p<0.05). Finally, we merged DMRs within 50 bp of one another.

For the F0 samples, we tested for both overall and sex-specific effects of hatchery rearing. For the overall effects of hatchery rearing, a generalized linear model (GLM) was used to test for the influence of origin on DNA methylation in the F0 while controlling for sex and the source by sex interaction. Based on the results and heatmaps for the F0, we split the hatchery parents into two cohorts: (i) seven “wildtype” fish whose DMR methylation patterns resembled wild fish, and (ii) nine “normal” hatchery fish whose DMR methylation patterns differed from wild fish (Figure S3). We consider epigenetic divergence between wild and hatchery-reared fish to be “normal” due to considerable evidence for the effects of hatchery rearing on the methylome of salmonids (e.g., reviewed in Koch et al. (2022)). We reran the DSS analysis for three separate F0 comparisons: comparing each hatchery subset independently to the wild fish and comparing the hatchery subsets to one another. We also split the full F0 dataset by sex and tested for sex-specific effects of hatchery rearing using a Wald test for the effect of origin for each sex, requiring a minimum delta (group difference in methylation) of 10% for both DMLs and DMRs. A GLM was used to test for maternal and paternal origin effects and their interaction in the F1. The pipeline is available at https://github.com/cvenney/dss_pipeline.

### Annotation and gene ontology enrichment analysis

Gene ontology annotation was performed with the DMR lists obtained from DSS. We annotated the genome using the GCF_905237065.1_Ssal_v3.1 transcripts file for the publicly available Atlantic salmon genome (https://www.ncbi.nlm.nih.gov/genome/?term=txid8030[orgn]) with GAWN v0.3.5 (https://github.com/enormandeau/gawn). We identified transcripts proximal to (±5 kb) the DMR positions from each DSS analysis and performed GO enrichment tests using the go_enrichment v1.1.0 pipeline (https://github.com/enormandeau/go_enrichment). The pipeline blasts the targets transcript against the Swiss-Prot database, obtains GO annotation information from the UniProt database, then uses GOATOOLS (Klopfenstein et al., 2018) to find significantly enriched GO terms.

## Results

We obtained an average of 301 153 191 raw methylation reads per sample and an average of 173 158 755 successful alignments. After all quality filtering, we retained a total of 25 451 589 CpG sites for the parental analysis and 31 440 242 for offspring (see Table S1 for detailed coverage information).

### Principal component and redundancy analyses

We analyzed 60 179 CpG sites with no missing data for F0 and 65 347 for F1 with PCA and RDA. We assessed methylation variation using PCA analysis and found whole genome methylation differences between hatchery reared and wild F0 salmon. The “wildtype” hatchery cohort identified in DMR analysis, as outlined in the methods, was intermediate between the wild and “normal” hatchery cohort (Figure 1A). Little differentiation between F1 groups was observed at the whole genome level (Figure 1B).

**Figure 1:**
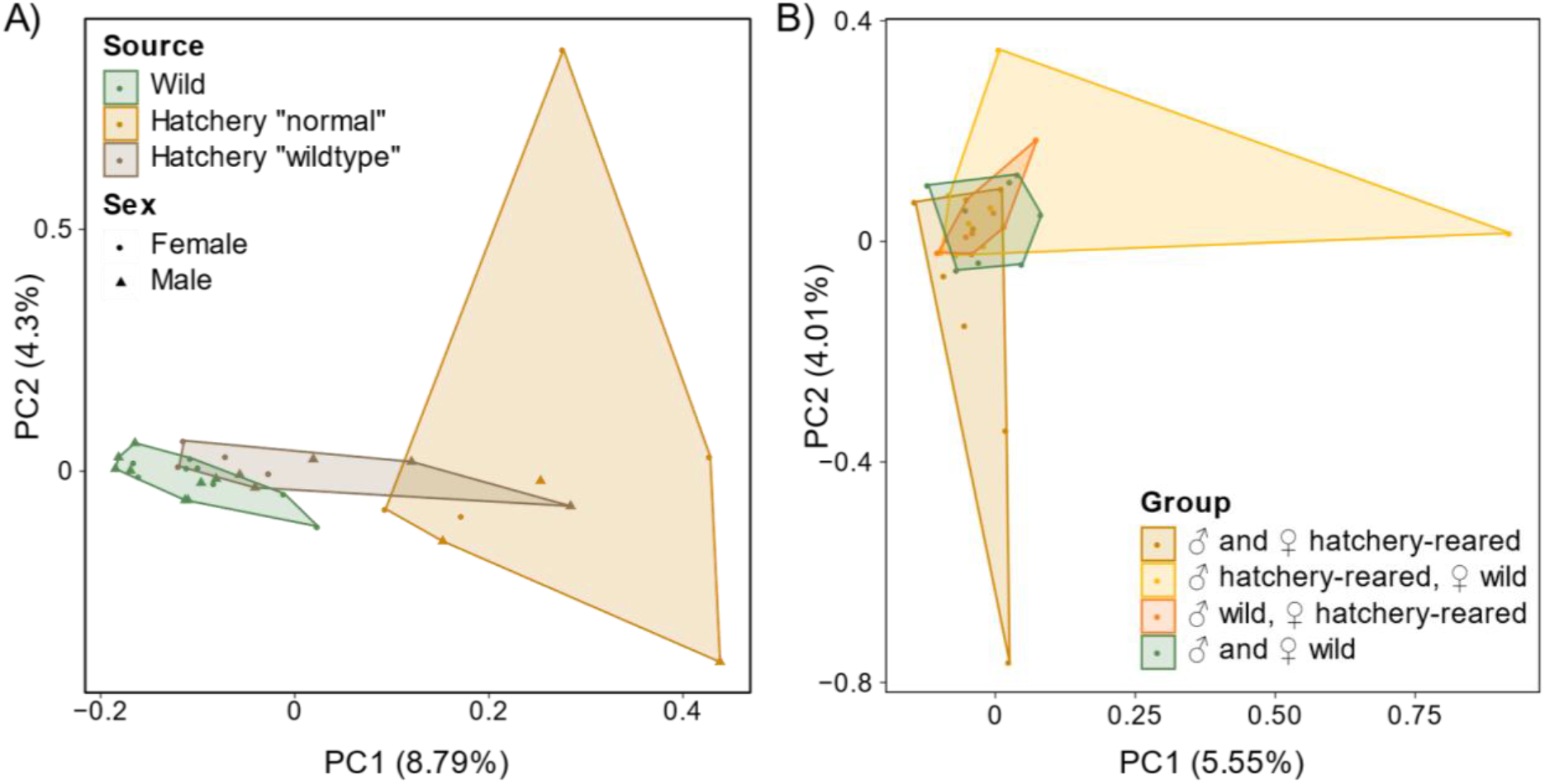
PCA plots for methylation data showing (A) differentiation between F0 hatchery cohorts and wild fish, and (B) lack of whole genome F1 differentiation based on parental rearing environment.

The F0 RDA model was significant (p=0.002, adjusted R^2^=0.023), as was the effect of source (p=0.001), though sex and the interaction effect were not significant (p=0.37 and p=0.26, respectively). The overall F1 RDA model was not significant (p=0.65), nor were the terms in the model (maternal source p=0.20, paternal source p=0.74, interaction p=0.87) indicating no significant whole genome effects of parental rearing environment.

### Differential methylation analysis: overall F0 effects of hatchery rearing

DSS analysis with all F0 samples identified two cohorts of hatchery fish, one “wildtype” cohort that clustered with wild fish and one “normal” hatchery cohort that differed from the wild and wildtype fish when controlling for sex and the source by sex interaction (Figure S3). For the full dataset, we found 165 DMLs and 266 DMRs due to origin. However, since there may be multiple age classes present in the hatchery fish and different manipulations have been performed on hatchery-reared fish by hatchery personnel, we split the F0 dataset into three groups: the two hatchery cohorts identified in the heatmap and PCA analysis, and the wild fish.

We identified 84 431 DMLs and 2 007 DMRs between wild and “normal” hatchery salmon (Figure 2A, Tables S2 and S3), with clear division between the two groups based on the dendrogram. GO enrichment analysis showed the DMRs were associated with organ morphogenesis, skeletal morphogenesis (e.g., GO:0035138 pectoral fin morphogenesis), and metabolic processes (Table S4).

**Figure 2:**
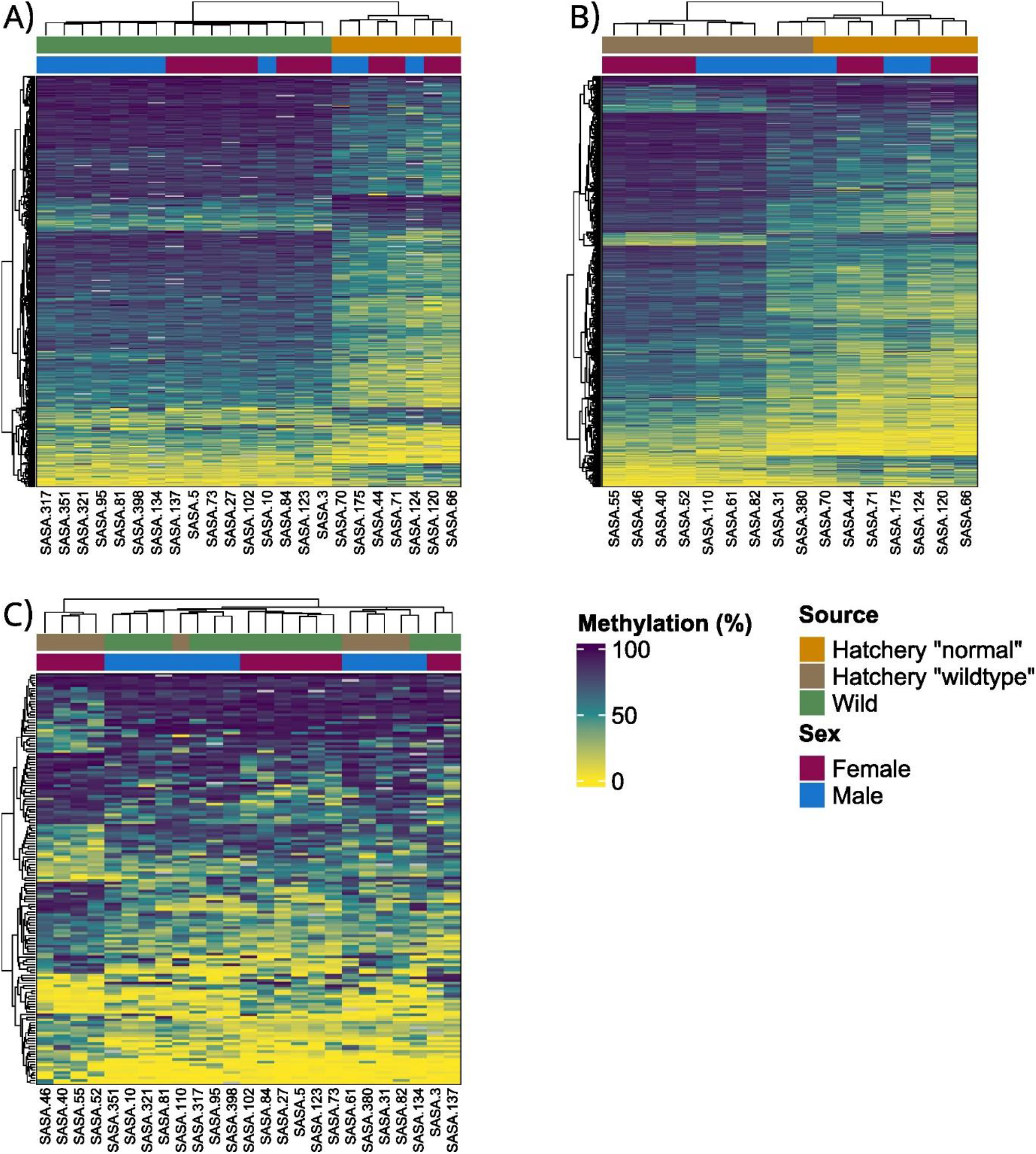
F0 differential methylation results between hatchery cohorts (“normal” = orange, “wildtype” = brown) and wild salmon (green). We detected (A) 2 007 DMRs between wild and “normal” hatchery salmon, (B) 1 285 DMRs between “normal” and “wildtype” hatchery cohorts, and (C) only 156 DMRs between the “wildtype” hatchery cohort and wild salmon. Hierarchical clustering was performed based on Euclidean distances. Percent methylation is denoted by the yellow to indigo scale for each DMR (0-100%).

We found 34 049 DMLs and 1 285 DMRs between “normal” and “wildtype” hatchery salmon (Figure 2B, Tables S5 and S6) with the two groups separating as predicted in the dendrogram other than SASA-31 and SASA-380, which were also the intermediate hatchery “wild type” points overlapping the hatchery “normal” polygon in Figure 1A. Enriched GO terms were associated with angiogenesis, intestinal morphology, skeletal morphology, and transcriptional processes (Table S7).

We only detected 84 DMLs and 156 DMRs between wild and “wildtype” hatchery salmon (Figure 2C, Tables S8 and S9) with no clear separation of groups in the heatmap, indicating that the “wildtype” hatchery cohort has a methylation profile similar to that of wild fish. Despite the small number of DMRs, there was functional enrichment for various transcriptional processes (Table S10).

### Differential methylation analysis: sex-specific F0 effects of hatchery rearing

We found considerable sex-specific effects of hatchery rearing in the F0. We detected 260 844 DMLs and 5 717 DMRs between male hatchery-reared and wild salmon (Figure 3A, Tables S11 and S12). We observed half the number of DMLs and DMRs due to hatchery rearing in female fish, with 100 451 DMLs and 2 439 DMRs due to source (Figure 3B, Tables S13 and S14), though both groups showed clear differentiation based on source in the heatmaps. The dendrograms also showed separate clustering of “normal” and “wildtype” female fish, though the cohorts were less differentiated in males (the “wildtype” samples SASA-31 and SASA-380 grouped with the “normal” hatchery males). We found only 734 overlaps between the male and female sex-specific analyses, indicating that most DMRs identified are sex-specific. There were 39 male-specific and 42 female-specific enriched GO terms with many overlaps between the sexes. Terms were associated with RNA synthesis and transcription for both sexes (e.g., GO:0010468 regulation of gene expression, GO:2001141 regulation of RNA biosynthetic process, GO:1903506 regulation of nucleic acid-templated transcription, GO:0006357 regulation of transcription by RNA polymerase II; Tables S15 and S16).

**Figure 3:**
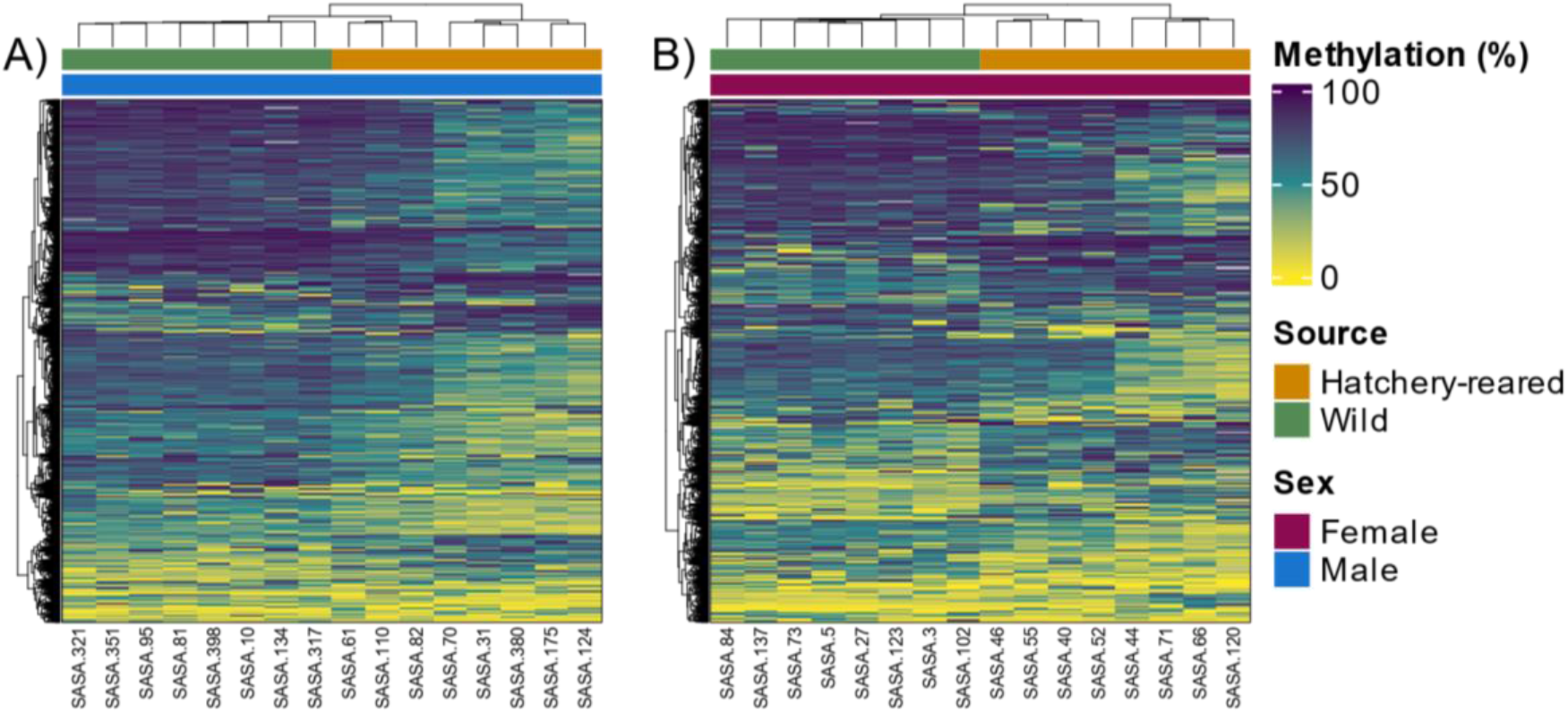
F0 sex specific DMRs due to rearing environment. We identified (A) 5 717 DMRs between male hatchery-reared and wild salmon, and (B) 2 439 DMRS between female salmon.

### Differential methylation analysis: F1 effects of parental hatchery rearing

The effects of hatchery rearing were diluted in the F1 fish compared to the F0. We observed 46 DMLs and 284 DMRs due to maternal source (Figure 4A, Tables S17 and S18), with clear division of offspring with wild dams and hatchery-reared sires, and hatchery-reared dams and hatchery-reared sires (offspring with wild sires were mixed regardless of maternal source). We found 11 DMLs and 254 DMRs due to paternal source (Figure 4B, Tables S19 and S20), with division occurring in offspring with hatchery dams based on paternal source, similar to the maternal DMRs. There were three DMLs and 271 DMRs due to the interaction between maternal and paternal source, with clear differentiation between ‘pure’ (two wild or two hatchery-reared parents) and hybrid offspring (Figure 4C, Tables S21 and S22). No GO terms were significantly enriched due to maternal source after filtering results. The paternal source DMRs showed enrichment for integral component of membrane (GO:0016021) and the interaction effect DMRs showed enrichment for regulation of cellular process (GO:0050794).

**Figure 4:**
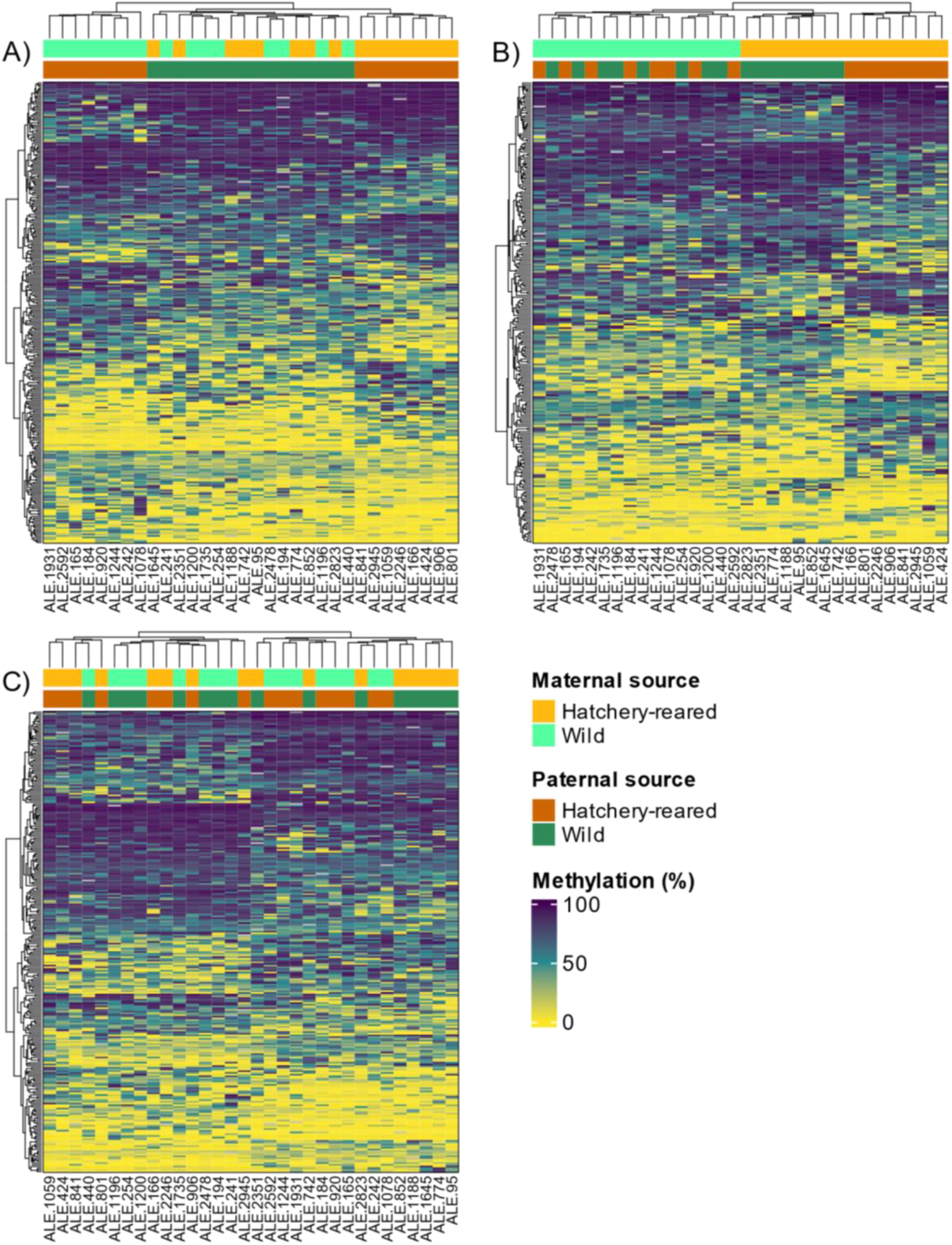
F1 DMR analysis identified (A) 284 DMRs due to maternal source, (B) 254 DMRs due to paternal source, and (C) 271 DMRs due to their interaction.

## Discussion

While the epigenetic effects of hatchery rearing in salmonids are well-documented (Gavery et al., 2019, 2018; Koch et al., 2022; Le Luyer et al., 2017; Leitwein et al., 2021, 2022; Rodriguez Barreto et al., 2019; Venney et al., 2021; Wellband et al., 2021), the long-term stability of these changes after release into natural systems has not been extensively studied, nor has the sex-specificity of the effects of hatchery rearing on the methylome. Here we showed that hatchery rearing led to general and sex-specific changes in Atlantic salmon fin DNA methylation that persisted until the salmon returned to spawn, consistent with similar results in Coho salmon returning to freshwater after eighteen months at sea (Leitwein et al., 2021). However, fewer methylation changes persisted to F1, indicating that the epigenetic effects of hatchery rearing primarily affected first-generation hatchery fish, though the fitness consequences of hatchery rearing may be persistent and insidious (O’Sullivan et al., 2020; Willoughby & Christie, 2019).

### Methylation analysis identified two hatchery cohorts

Two cohorts of hatchery salmon were identified through both PCA for whole genome methylation and DMR analysis (Figures 1A, 2A-C, S2) with the “wildtype” cohort more epigenetically similar to wild fish than the “normal” hatchery cohort. The latter was named “normal” due to the expectation that hatchery rearing influences the methylome based on the current literature (e.g., reviewed in Koch et al., 2022). The cohorts were not related to sex or time spent at sea (one winter vs. multiple winters at sea) and there were no particular rearing manipulations performed on the F0 during their time in the hatchery. However, hatchery origin fish were released at different locations down the Rimouski River. Therefore, they could have experienced different conditions during the final stage of transportation and stocking (e.g., due to temperature). Each year, some parr were released in the same areas as wild conspecifics whereas others were stocked in a section of river upstream of a waterfall where wild salmon parr are not found. Regrettably, we could not determine the release site of each fish as they were not tagged or otherwise identifiable after release. Thus, it is possible that the cohorts exhibit different methylation patterns due to differences in early life environment after stocking, with “wildtype” hatchery salmon assuming methylation patterns similar to wild fish due to a shared environment. It is also possible that hatchery rearing had a negligible effect on methylation in our study and the epigenetic differences between groups are due to differences in the environment where they were released. However, even the “wildtype” hatchery fish show methylation differences compared to wild fish, and we observed epigenetic inheritance of these effects in F1 regardless of parental cohort, indicating that the hatchery environment does influence the methylome. Indeed, hatchery rearing has been shown to affect the methylome even after oceanic migration in Coho salmon (Leitwein et al., 2021). However, it is possible that the residual effects of hatchery rearing were diminished due to the shared river environment after release. It is also possible that other tissues would retain the methylation changes associated with hatchery rearing, though we used fin in our study to allow nonlethal sampling of F0 during their spawning migration.

The functional significance of DMRs between the two hatchery cohorts was consistent with previously reported phenotypic effects of hatchery rearing, further suggesting that the methylation differences were due to the hatchery environment rather than the river environment. GO enrichment analysis showed that the “normal” and “wildtype” hatchery fish exhibited methylation differences at regions associated with angiogenesis, intestinal morphology, skeletal morphology, and transcriptional processes. A previous study comparing mangrove killifish (*Kryptolebias marmoratus*) reared in both barren tanks and environments enriched with logs and plants reported methylation changes in genes involved in angiogenesis (Berbel-Filho, Rodríguez-Barreto, Berry, Garcia De Leaniz, & Consuegra, 2019). Another study using Rimouski salmon showed that hatchery rearing alters the gut microbiota (Lavoie et al., 2018) which may be associated with morphological changes (Liu, Wang, Ou, Hou, & He, 2020). Thus, the two hatchery cohorts may have diverged due to differences in early life environment with the “normal” cohort harbouring methylation changes associated with the typical phenotypic effects of hatchery rearing.

Comparisons between wild and “normal” hatchery fish DMRs also showed functional enrichment for organ and skeletal morphogenesis including pectoral fin morphogenesis, and metabolic processes. Hatchery-reared Atlantic salmon have lower swimming performance relative to wild conspecifics (Pedersen et al., 2008). Thus, differences in fin DNA methylation between “normal” hatchery fish and wild fish are associated with biological functions that may underly previously reported differences in swimming capacity, though further studies in other tissues are needed for a mechanistic link.

In contrast, “wildtype” hatchery fish and wild fish were epigenetically similar relative to other comparisons, though methylation differences were associated with transcriptional regulation. Therefore, the epigenetic effects of hatchery rearing could potentially be minimized through supplementation efforts. We suspect this is due to the “wildtype” hatchery fish sharing an environment with wild fish after stocking, though regrettably we cannot definitively determine the causative factor(s) leading to epigenetic similarities between “wildtype” hatchery and wild fish. Further experimental investigation would be needed to determine how to minimize the epigenetic effects of hatchery rearing. Overall, we show that the epigenetic effects of early rearing environment are not short-term acclimatory responses, but rather lead to long-term effects on the methylation states of salmon that persist until spawning (i.e., after two to three years living in rivers followed by one to two years spent at sea).

### Sex-specific effects of hatchery rearing

We provide evidence that hatchery rearing affects the epigenome in a sex-specific way, further complicating our understanding of the effects of hatchery rearing on the methylome. We identified 5 717 male-specific and 2 439 female-specific F0 DMRs due to rearing environment, with only 734 DMRs shared between the sexes. Sex-specific methylation differences have been shown to occur in fish (Fellous et al., 2018; Laing et al., 2018; Podgorniak, Brockmann, Konstantinidis, & Fernandes, 2019). In Nile tilapia (*Oreochromis niloticus*), early domestication (i.e., wild-born fish transferred to a hatchery for domestication) led to sex-specific methylation responses to the hatchery environment, with only 57 of ~1000 DMLs shared between male and female fish (Podgorniak et al., 2019). Transcriptional differences between the sexes have been reported in brook charr (*Salvelinus fontinalis*; Sutherland, Prokkola, Audet, & Bernatchez, 2019) and are associated with DNA methylation differences in zebrafish (*Danio rerio*; Laing et al., 2018). While hatchery fish generally have lower reproductive success than wild conspecifics regardless of sex (Bouchard et al., 2022; Christie et al., 2014; O’Sullivan et al., 2020), a previous meta-analysis showed that hatchery rearing had a greater effect on the reproductive success of males than females when compared to wild fish of the same sex (Christie et al., 2014). Female hatchery fish tend to have higher reproductive success in the wild than male hatchery fish (Berntson, Carmichael, Flesher, Ward, & Moran, 2011; Christie et al., 2014; Milot et al., 2013; Thériault, Moyer, Jackson, Blouin, & Banks, 2011; Williamson, Murdoch, Pearsons, Ward, & Ford, 2010) and thus molecular responses to hatchery rearing are likely complicated by sex. While the genomic regions affected by hatchery rearing differ between sexes, GO enrichment analysis showed that the regions were associated with many of the same processes in both males and females, with 35 of 39 enriched male-specific GO terms also present in the female analysis. Due to the potential for epigenetic inheritance, sex-specific epigenetic effects could also lead to parental rearing environment affecting offspring epigenetics in complex ways based on which parent(s) experience the hatchery environment, and whether maternal or paternal effects occur in specific regions of the genome.

### Dilution of epigenetic effects of hatchery rearing in F1

Despite the persistent effects of hatchery rearing in F0, we observed relatively little inheritance to F1 based on parental environment: there were few F1 DMLs and DMRs based on parental source (Figure 4A-B) and there was no evidence for genome-wide methylome differentiation (Figure 1B). While there is evidence for epigenetic inheritance across taxa, it is unclear to what extent the epigenome is inherited, though there is often a dilution of inherited effects over generations (Anastasiadi et al., 2021). Studies have shown that hatchery rearing can influence sperm methylation (Gavery et al., 2018; Leitwein et al., 2021; Wellband et al., 2021), though somatic methylation changes are often not present in the F0 germline (Anastasiadi et al., 2021; Gavery et al., 2019, 2018) and the genomic regions showing differential methylation often differ between F0 sperm and F1 somatic tissues (Wellband et al., 2021). Thus, despite considerable epigenetic effects of hatchery rearing in F0, many of these changes are not passed on to F1.

The few methylation differences that were inherited based on parental rearing environment exhibited complex, nonadditive effects that may be difficult to account for during conservation efforts. When testing for maternal influences on offspring methylation, differences were primarily between offspring from wild and hatchery dams with hatchery-reared sires, whereas offspring from wild sires were similar regardless of maternal origin (Figure 4A). The same effect was observed when testing for paternal influences, with paternal rearing environment mainly affecting offspring methylation when the dams were of hatchery origin (Figure 4B). This suggests nonadditive epigenetic inheritance based on hatchery rearing wherein the origin of both parents influences the DNA methylation state of offspring, rather than simple maternal or paternal effects. A study in Chinook salmon (*Oncorhynchus tshawytscha*) showed that hatchery-origin fish had greater nonadditive effects on DNA methylation when reared in a semi-natural environment as opposed to a hatchery environment (Venney et al., 2021), though a study on reciprocal hybrid steelhead (*O. mykiss*; hatchery dam and wild sire, or wild dam and hatchery sire) identified additive, heritable effects on gene expression (Christie et al., 2016). Thus, is it possible that the complexity of the natural environment led to complicated epistatic effects resulting in nonadditive effects on methylation, though the link between DNA methylation and gene expression may be complex (Christensen et al., 2021).

Unexpectedly, the maternal by paternal interaction effect DMRs showed two clusters: the pure strain (hatchery-hatchery and wild-wild) and hybrid offspring. While hybridization can often cause atypical methylation patterns compared to pure strains (Gimenez, Vazquez, Trepat, Cambiaso, & Rodríguez, 2021; Raza et al., 2017; Shen et al., 2012), it was surprising that pure wild and pure hatchery-origin offspring had similar methylation patterns at these loci. It is possible that hybridization between wild and hatchery salmon led to dysregulation of epigenetic marks, though we find this an unlikely explanation after a single generation of hatchery rearing.

Overall, these complex nonadditive patterns of inheritance based on parental environment have the potential to influence offspring phenotype and fitness. Indeed, parental hatchery rearing has been associated with fitness-related phenotypic variation in F1 in Atlantic salmon (Wellband et al., 2021), though F1 effects of parental hatchery rearing are complex and weaker than F0 effects in our study. It is possible that slight differences in offspring environment could contribute to the complex patterns of inheritance we observed. The timing of juvenile release can also influence the fitness consequences of hatchery rearing (Milot et al., 2013), therefore the timing of F0 release may influence the extent of epigenetic inheritance due to hatchery rearing and merits further study. Due to the nonadditive nature of these heritable effects, it may be difficult to predict epigenetic responses to hatchery rearing beyond F0 and ensure effective conservation actions, though we come to the encouraging conclusion that the epigenetic effects of hatchery rearing are clearly diluted in F1 offspring.

### Conclusions

We showed that fin DNA methylation changes associated with hatchery rearing persisted in Atlantic salmon after release into natural systems and until spawning. The detection of two cohorts of hatchery fish suggests that slightly modified supplementation techniques may be able to reduce the epigenetic effects of hatchery rearing, though further research is needed to determine what factors may decrease the extent of these differences. In particular, future studies attempting to minimize the epigenetic effects of hatchery rearing will need to characterize the methylome of several different tissues in order to determine the overall systemic implications of manipulated environments on the methylome once the cost of whole genome methylation sequencing declines. We also observed sex-specific effects of hatchery rearing in F0 which may contribute to fitness differences between male and female hatchery-reared fish (Christie et al., 2014). We showed that few of these effects persisted to F1, suggesting that the effects of hatchery rearing may disappear rapidly over generations. However, the heritable effects of hatchery rearing exhibit complex, nonadditive patterns dependent on both maternal and paternal origin which may affect phenotype and will be difficult to account for in conservation and management decisions. With increasing investment and research into improving hatchery rearing and supplementation efforts, understanding and minimizing the detrimental effects of supplementation on fish is critical to ensure effective conservation efforts.

## Acknowledgements

We appreciate Dr Mark Christie’s helpful input and discussion of our results. We thank Gabriel Piette-Lauzière, Charles Babin, and Alexandre Carbonneau for assistance in the laboratory and sample preparation. We also thank Subject Editor Dr. Luciano Beheregaray and the anonymous reviewer for their help in improving the manuscript. This study was funded by the Ministère des Forêts, de la Faune et des Parcs du Québec (MFFP) and by NSERC (RDCPJ 537349–18).

## Data Accessibility and Benefit-Sharing

Raw whole genome methylation data and metadata are deposited in the SRA (BioProject PRJNA892473) and will be released publicly upon acceptance. Benefit-sharing statements are not applicable.

## Author Contributions

CJV, LB, JA, GC, and RB conceived of the project. CJV, RB, and LL performed the analyses. CJV wrote the initial manuscript and all authors contributed to the final version. Funding was provided by the Ministère des Forêts, de la Faune et des Parcs du Québec (MFFP) and an NSERC grant to LB (RDCPJ 537349 – 18).

