## Supplementary Figures S1-S2 for "Captive rearing effects on the methylome of Atlantic salmon after oceanic migration: sex-specificity and intergenerational stability"

**Figure S1:** Experimental design for sampling of Rimouski salmon F0 and F1 for methylation analysis.

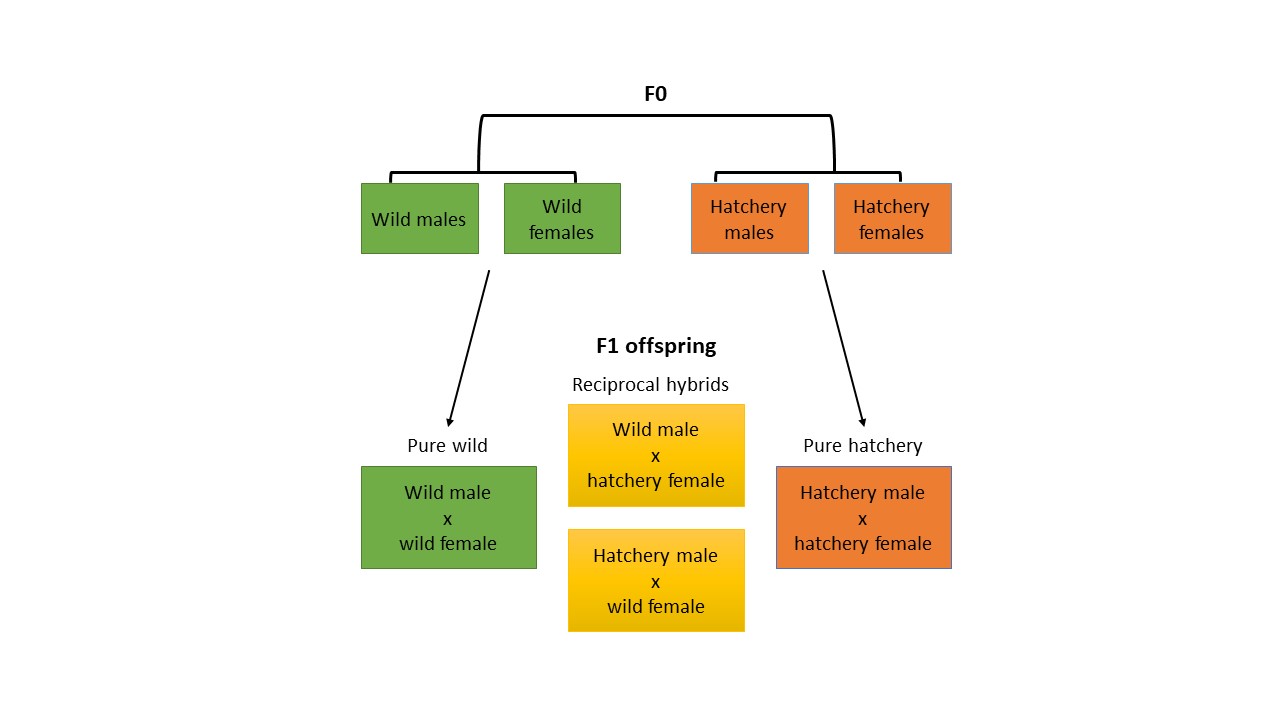

**Figure S2:** Principal component analysis based on 33 microsatellite loci for 30 samples per population (N=90) with maximum 5% missing data. Weir and Cockerham’s pairwise F_ST_ was also calculated as follows for each pair: Rimouski-Puyjalon: 0.098, Rimouski-Romaine: 0.104, Puyjalon-Romaine: 0.125.

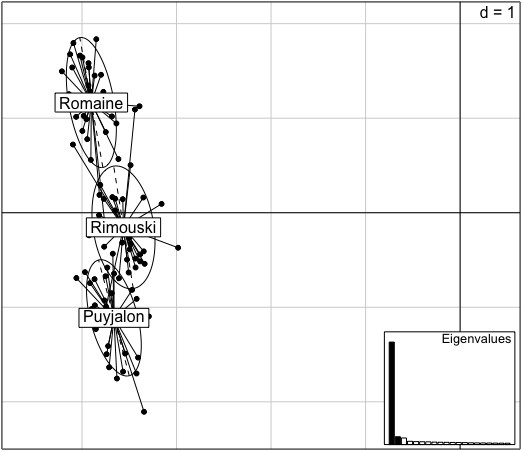

**Figure S3:** Overall F1 DMR based on source (hatchery-reared or wild). The dendrogram based on Euclidean distance along the top of the graph separated the wild salmon by sex but identified two cohorts of hatchery-reared salmon which were then analyzed as separate hatchery cohorts (“wildtype” and “normal”).

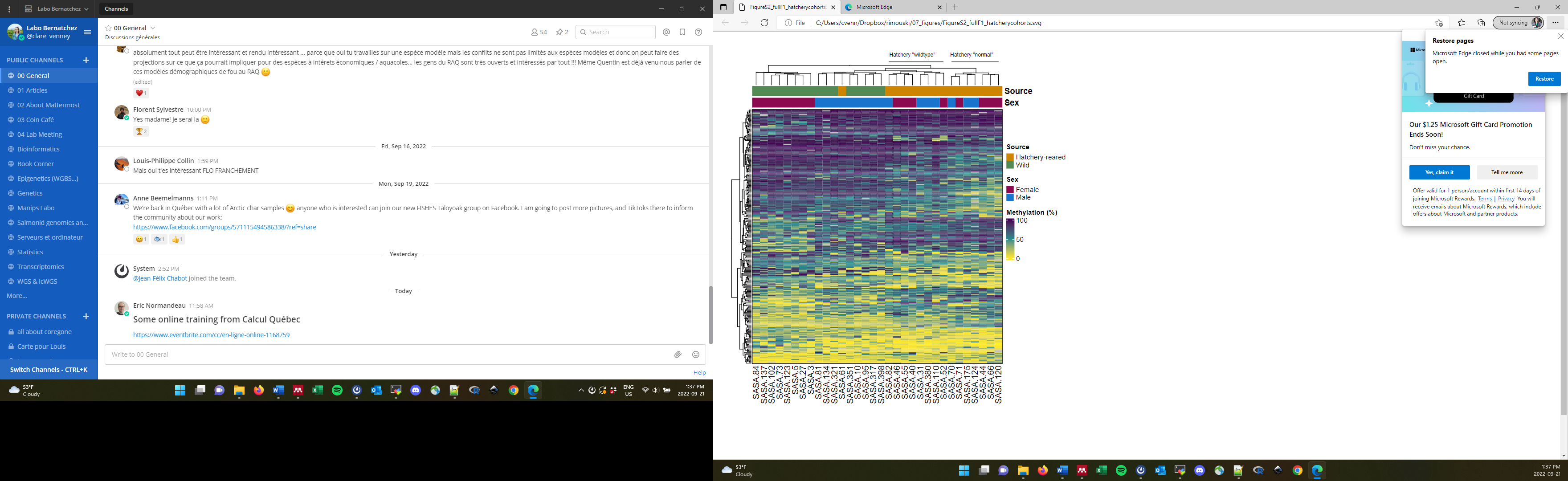
